# Limiting intestinal iron absorption rescues glial defects and extends lifespan in a Drosophila model of Friedreich’s ataxia

**DOI:** 10.64898/2026.06.04.730074

**Authors:** Ema Turki, Estelle Jullian, Pierre Delamotte, Anne Filipe, Laura Tixier-Cardoso, Sandrine Middendorp, Elodie Martin, Véronique Monnier

## Abstract

Friedreich ataxia (FRDA) is a neurodegenerative and cardiac disease caused by GAA repeat expansions within the first intron of the *FXN* gene, leading to reduced frataxin expression. Frataxin is required for iron sulfur cluster (ISC) biosynthesis, and its deficiency results in multiple cellular dysfunctions, including mitochondrial iron overload. Although altered iron homeostasis has been reported in several frataxin-deficient models and in FRDA patients, its contribution to disease progression remains debated. Here, we used a GAA expansion-based *Drosophila* model of FRDA, termed fh-GAAs, to investigate the impact of reducing intestinal iron absorption on disease progression. We first found that iron accumulation was tissue-specific and predominantly affected the central nervous system. Furthermore, glial cells were affected more severely than neurons, suggesting an increased vulnerability of glia to frataxin deficiency. Reducing intestinal iron uptake, either through treatment with bathophenanthroline disulfonic acid (BPS), an extracellular iron chelator, or by gut-specific silencing of the iron transporter *Malvolio*, nearly doubled fly survival. BPS treatment also improved sensitivity to dietary iron, enhanced locomotor performance, fully restored normal brain size, and prevented glial alterations. Altogether, our findings identify glial cells as early and preferential targets of frataxin deficiency in an iron-dependent manner and support the *in vivo* relevance of intestinal iron uptake as a potential modulator of disease severity in FRDA.

## 1. Introduction

Friedreich Ataxia (FRDA) is a progressive autosomal recessive neurodegenerative disease characterized by gait and limb ataxia, muscle weakness, loss of proprioception and dysarthria (Durr et al., 1996; Harding, 1981; Pandolfo, 2009). Beyond neurological impairment, patients also develop hypertrophic cardiomyopathy, which constitutes the main cause of death (Tsou et al., 2011), diabetes and scoliosis. FRDA is caused by a GAA trinucleotide repeat expansions within the first intron of the *FXN* gene leading to decreased gene expression (Campuzano et al., 1996). *FXN* encodes frataxin, a mitochondrial protein involved in the biosynthesis of iron-sulfer clusters (ISC), which are essential cofactors for a large number of proteins (Braymer and Lill, 2017; Gervason et al., 2019). Frataxin deficiency leads to multiple cellular dysfunctions that have been described in patients and models of the disease, most notably dysfunctions of the mitochondrial respiratory chain, perturbations of iron and lipid homeostasis, and hypersensitivity to oxidative stress (Llorens et al., 2019; Tamarit et al., 2016). Mitochondrial iron accumulation has been observed in multiple frataxin-deficient cellular and animal models, as well as in the heart and nervous system of patients (La Rosa et al., 2020). However, the precise mechanisms involved in this iron accumulation as well as its contribution in the progression of the disease remain subjects of debate. Therapeutic strategies targeting iron overload, such as the use of deferiprone, have been explored. Deferiprone is an iron chelator able to cross the blood-brain barrier and cellular membranes and to shuttle iron between subcellular compartments (Sohn et al., 2008). While it was proved protective in cellular and animal models, (Kakhlon et al., 2008; Soriano et al., 2013; Whitnall et al., 2008), it led to inconclusive results when it comes to clinical trials, with safety concerns at high doses (Pandolfo et al., 2014; Pandolfo and Hausmann, 2013). Recently, attention has shifted toward ferroptosis, a regulated form of non-apoptotic cell death driven by iron-dependent accumulation of lipid peroxides with compromised glutathione-dependent repair systems (Dixon et al., 2012). Frataxin-deficient cells exhibit hypersensitivity to the ferroptosis-inducer erastin (Cotticelli et al., 2019; Du et al., 2020). In addition, hallmarks of ferroptosis have been described in several frataxin-deficient cells, including mouse dorsal root ganglia (DRG) neurons and cardiomyocytes as well as patient-derived fibroblasts (La Rosa et al., 2021; Sanz-Alcazar et al., 2024). These findings reinforce the rationale for targeting iron-dependent damage as a therapeutic strategy. In this study, we aimed to investigate the impact of limiting intestinal iron uptake on disease progression. Frataxin is a highly conserved mitochondrial protein, making Drosophila a powerful *in vivo* model for FRDA and a highly relevant model to address this issue (Llorens et al., 2019). We previously generated Drosophila models in which GAA expansions have been inserted in the intron of the frataxin homolog gene *fh* (Jullian et al., 2024; Russi et al., 2020). Specifically, the Drosophila flies carrying a 42 GAA expansion (fh-GAAs), exhibit a strong decrease in fh expression, a shortened lifespan, progressive locomotor dysfunctions, hypersensitivity to oxidative stress and cardiac dilatation (Jullian et al., 2024). These GAA-expansion models have proven valuable for drug screening, transcriptional profiling, suppressor gene identification and contributed recently to the discovery of a novel regulatory mechanism of iron-sulfur cluster biosynthesis (Jullian et al., 2024; Russi et al., 2020; Want et al., 2025). Here, we first assessed the extent of iron dyshomeostasis in fh-GAAs flies. We then tested whether limiting iron availability could modify disease progression using two complementary approaches : pharmacological reduction of intestinal iron uptake with the BPS extracellular chelator, and gut-specific RNAi-mediated knockdown of the iron transporter Malvolio. Both strategies markedly improved adult survival. BPS treatment also improved adult locomotor performance but not cardiac dilatation. Histological analysis of the central nervous system of fh-GAAs flies revealed a predominant impact on glial cells in the larval ventral nerve cord compared with neurons and the altered glial nuclear morphology was fully prevented by BPS treatment. Overall, these results identify glial cells as early iron-sensitive targets of frataxin deficiency and support a beneficial role of modulating dietary iron availability in a Drosophila model of FRDA.

## 2. Materials and methods

### 2.1 Drosophila lines, culture medium and drug treatments

The fh-GAAs line was described in (Jullian et al., 2024). *fh-GAAs/FM7i* females were regularly crossed with *FM7i/Y* males of the *AMPKalpha3/FM7i* line to prevent autosomal genetic drifts of the line. Every five to ten outcrosses, corresponding to every four to six months, the line is amplified and used for the experiments of the following four to six months. The *Mex-GAL4* ; *UAS-dicer2* line was kindly provided by Jacques Montagne. The *w*^*1118*^ strain, used as a control line, and the Malvolio-RNAi line (v109434) were obtained from the Vienna Drosophila Resource Center (VDRC). The *AMPKalpha3/FM7i* line was obtained from the Bloomington Stock Center. The standard fly medium contained 82.5 mg/ml yeast, 34 mg/ml corn meal, 50 mg/ml sucrose, 11.5 mg/ml agar, and 27.8 μl/ml methyl 4-hydroxybenzoate (stock solution 200 g/l in ethanol). For compound incorporation, standard agar was replaced by Low Melting Agarose (14mg/ml, Merck REF.A9414) and the medium was equilibrated at 37°C prior to drug incorporation to prevent thermal degradation. Deferiprone (Merck REF.Y0001976, stock solution 100mM in H_2_0), deferoxamine mesylate (Merck REF.D9533, 100 mM in H_2_0) and BPS (Merck REF.146617, 10 mM in H_2_0) were incorporated from stock solutions in the food at indicated final concentrations. When required, RU486 (Betapharma) was incorporated from a 20 mg/ml stock solution in ethanol, or a similar amount of ethanol for untreated controls.

### 2.2 Lifespan analysis and spontaneous locomotor activity

Adult male flies were collected within 24 h of eclosion, housed in tubes by groups of 30, and raised at 26 °C under a 12 h–12 h light-dark cycle. For lifespan experiments, the tubes were changed every two days to provide fresh food, and dead flies were counted. For locomotor activity, 3 days old males were individually placed in glass tubes containing medium composed of 5% sucrose and 1.35% agar. Activity was recorded with the Drosophila Activity Monitor single-beam (DAM) system (TriKinetics, Waltham, MA) and the data were analyzed with the ShinyR-DAM software (Cichewicz and Hirsh, 2018).

### 2.3 Viability measurements

Ferric ammonium citrate (Merck REF.F5879) and copper sulfate (Merck REF.451657), with or without BPS, were incorporated in the medium on which *fh-GAAs/FM7i* females crossed with *w*^*1118*^ males were allowed to lay eggs. Adult progeny were counted (5 to 10 tubes per condition for copper sensitivity, 10 tubes per condition for iron sensitivity) and the number of *fh-GAAs/+* and *FM7i/+* females were used to estimate the expected number and viability of *fh-GAAs/Y* males.

### 2.4 Ferrozine assays

The protocol was adapted from (Missirlis et al., 2006). Thirty third instar male larvae per sample were thoroughly grinded with a pestle in 150 µl of RIPA1X (BIO BASIC CANADA INC. REF :RB4477) on ice. Samples were centrifuged at 16,000xg for 20 minutes at 4°C and supernatants were collected. HCl was added to a final concentration of 3.7%, and samples were heated at 95°C for 20 minutes. Tubes were let 5 minutes at room temperature before being centrifuged at 16,000xg for 20 minutes at room temperature. In a transparent multi-well plate, was added in this order: 50 µL of supernatant, 20 µL of 75 mM ascorbate (Merck REF :11140), 40 µL of 10M ammonium acetate (Merck REF :A7262) and 20 µL of 10 mM ferrozine (Merck REF :160601). The plate was sealed with a plastic foil and packed in aluminum foil, gently shaked for 10 minutes at room temperature. Absorbance was read at 562nm using a plate reader (FlexStation 3R Molecular Devices).

### 2.5 Iron staining and CNS size measurements

To detect ferric iron, third instar male wandering larvae were dissected in PBS and quickly transferred to 3,7 % formaldehyde in PBS for 15 min. Following 3 washes of 5 min in PBS 0,4 % Triton, the samples were incubated in the dark in the working iron solution (Iron Stain Kit, Merck REF :HT20) for 90 min. After 3 washes of 5 min in PBS 0,4 % Triton, they were incubated 2 hours in a DAB solution (Dab Substrate, Roche REF : 11718096001). Samples were washed 3 times in PBS 0,4 % Triton and mounted on glass slides. For larval CNS measurements, brains were dissected in PBS, fixed for 30 minutes in formaldehyde 3.7%/PBS, washed 3 times in PBS, and mounted on glass slides in ProLong Diamond Antifade Mountant. Images were taken onto a Zeiss Axio Observer Z.1 microscope and processed using Fiji.

### 2.6 Immunofluorescence staining

Third instar male wandering larvae were dissected in PBS and fixed for 30 min in 3.7% formaldehyde at room temperature (RT). Incubations with primary antibodies were performed overnight at 4°C and with secondary antibodies at RT for 2 hours in PBST (0.5% Tween-20 in 1X PBS) supplemented with 2% BSA. The following concentrations were used : mouse anti-Repo (8D12, DSHB), 1 :40 ; rat anti-Elav (7E8A10, DSHB) 1:40 dilution ; goat anti-mouse Alexa Fluor Plus 488 (A32723, Invitrogen), 1 :500 ; goat anti-rat Alexa Fluor Plus 647 (A48265TR, Invitrogen), 1:500. Larval brains and nerve cords were mounted in ProLong Diamond Antifade Mountant. Confocal imaging was performed using a LSM 980 Airyscan 2 FLIM with a 63X objective. Nuclei aspect ratio and size were quantified on comparable z-stacks using the Analyze Particles function in Fiji/ImageJ. The aspect ratio was defined as the ratio between the longest and the shortest nuclei axis. Neuronal and glial nuclei were manually counted and annotated using the Multi-point Tool in Fiji.

### 2.7 Cardiac imaging

To monitor cardiac function, *fh-GAAs* or *w*^*1118*^ females were crossed with *HandGS>mitoGFP* males. In the progeny, *fh-GAAs/Y* ; *HandGS>mitoGFP* and control male flies were collected and treated with 100 µg/ml RU486, leading to the expression of a mitochondrial targeted GFP specifically in cardiomyocytes. This allowed the labelling of the heart and the recording of cardiac beats in vivo on flies anaesthetized with FlyNap (Sordalab). Recording of the movies and extraction of cardiac parameters were performed as previously described in (Monnier et al., 2012).

### 2.8 Quantification of transcripts by qRT-PCR

Total RNA were extracted as described in (Reinhardt et al., 2012). Samples were treated with dsDNase (Thermo Fisher Scientific) according to the manufacturer’s instructions. cDNAs were synthetised from isolated total RNA samples using SusperScript™III Reverse Transcriptase (Thermo Fisher Scientific). qPCRs were performed with the qPCR Mix (Promega) on a LightCycler480 (Roche). The ribosomal gene rp49 was used as an internal reference for normalization. The sequences of primers are provided in supplementary Table S1. Standard curves were performed to check the efficiency of the primers. The ΔΔCt method was used for relative quantifications that were made on four independent biological samples with four technical replicates for each condition.

### 2.9 Statistical analysis

All statistical tests were assessed using Prism software (GraphPad, PrismV11.0.0). Individual values are shown as dots with mean and standard deviation.

## 3. Results

### 3.1 Iron accumulates in the central nervous system of fh-GAAs frataxin-deficient flies

We first aimed to assess iron metabolism in frataxin-deficient flies. Since *fh* is located on the X chromosome, all experiments were performed on hemizygous male flies carrying the fh-GAAs allele. We quantified on male third instar larvae the expression levels of genes encoding the major regulators of iron homeostasis : Malvolio (Mvl), the Drosophila homolog of DMT1 involved in iron intestinal absorption, the homologs of ferritin 1 Heavy Chain (Fer1HCH) and ferritin 2 light chain (Fer2LCL) involved in iron storage and absorption, the transferrin Tsf1, abundant in hemolymph and potentially involved in iron transport and finally mitoferrin (Mfrn) involved in mitochondrial iron import (reviewed in (Mandilaras et al., 2013). We did not detect significant systemic differences in expression levels of these genes in *w*^*1118*^ control and fh-GAAs third instar larvae, except for *Mfrn* that was slightly decreased (Fig. 1A). We also quantified total iron levels that were unchanged in frataxin-deficient larvae compared to controls in ferrozine assays (Fig. 1B). Next, we performed Prussian blue staining, that detects Fe^3+^, on specific tissues. We observed a strong iron accumulation in the larval brain and the neuropile of the ventral nerve cord in fh-GAAs mutants (Fig. 1C). Fe^3+^ accumulation was also observed compared to controls in pericardial cells (Fig. 1D) but not detectable in other tissues, such as the heart (Fig. 1D), the gut (Fig. 1E) or imaginal discs (Fig. 1F). Consequently, Fe^3+^ accumulation appears to be tissue-specific and affects mainly the central nervous system of frataxin-deficient flies.

**Figure 1:**
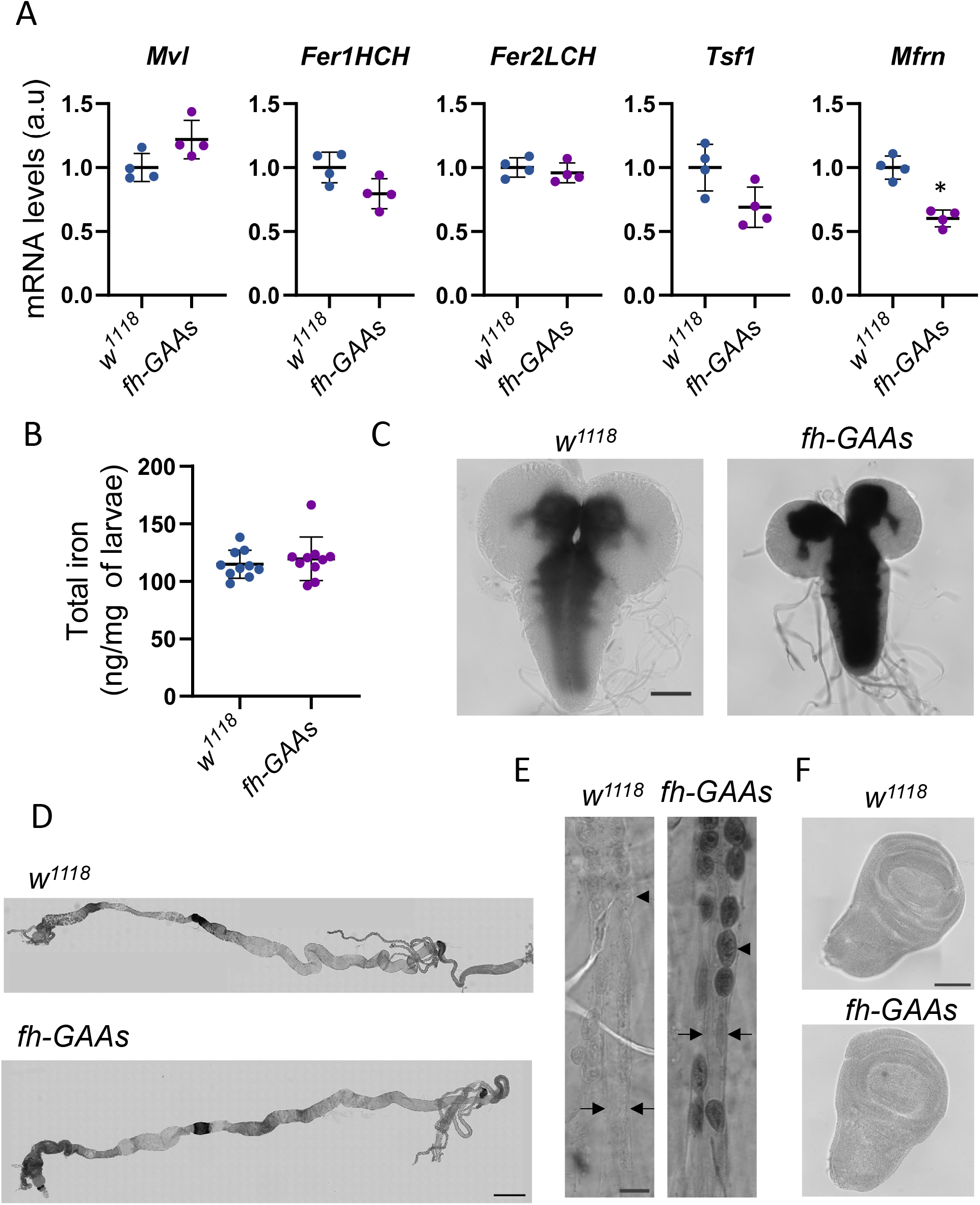
Expression of iron-regulatory factors and iron accumulation in frataxin deficient flies. A. Quantification of *Fer1HCH (Ferritin 1 heavy chain homologue), Fer2LCL (Ferritin 2 light chain homologue), Tsf1 (*Transferrin 1, *Mfrn* and *Mvl* transcript levels in control *w*^*1118*^ and *fh-GAAs* male third instar larvae. Significant difference is indicated : * P<0.05 (two-sided non-parametric Mann-Whitney tests). Dots correspond to independent biological samples each obtained from 20 individuals. B. Quantification of total iron levels with ferrozine assays. Dots correspond to independent biological samples each obtained from 30 individuals. C-E Prussian blue-DAB staining of larval *w*^*1118*^ and fh-GAAs tissues. C : larval brains (scale bar : 100 µm), D : Guts (scale bar : 500 µm). E : Heart and pericardial cells (scale bar : 50 µm, arrows indicate lateral sides of the cardiac tube, arrow heads the pericardial cells). F : Wing imaginal discs (scale bar : 100 µm).

### 3.2 BPS-feeding and gut-specific down regulation of the Malvolio iron transporter improves survival of frataxin-deficient flies

Following these observations, we investigated whether treating frataxin-deficient flies with iron chelators could improve their survival. As previously reported, fh-GAAs male flies are viable but exhibit a severely reduced lifespan, with a mean of less than 10 days compared to 40 days in control flies (Jullian et al., 2024). We first tested deferiprone, which failed to improve fh-GAAs survival (Fig. 2A). This lack of efficiency contrasts with a previous study in another RNAi-based FRDA *Drosophila* model (Soriano et al., 2013), suggesting a model- or dosage-dependent efficiency. Treatment with 100 µM deferoxamine mesylate significantly extended both median and mean lifespan by 21% and 23%, respectively (Fig. 2B). The most robust protection was achieved with 100 µM BPS, which increased median and mean lifespan by 80% and 93%, respectively (Fig. 2C). Notably, this increased survival was observed when the treatment was administered during the developmental stage. Treatment during the adult stage provided no additional survival advantage (Fig. S1A). Furthermore, increasing the BPS concentration to 250 µM was less effective than the 100 µM dose (Fig. S1B). This indicates that the protective effects of BPS are stage- and dose-dependent. Next, we quantified *fh* transcript levels in fh-GAAs and *w*^*1118*^ control flies, treated or untreated with BPS. The treatment did not modify *fh* expression, ruling out any protective effect linked to increased expression (Fig. S1C). BPS is a membrane-impermeable iron chelator known to reduce intestinal iron absorption in flies (Hernandez-Gallardo et al., 2024; Mehta et al., 2009). To confirm that reducing intestinal iron absorption was protective, we aimed to use an independent approach. Malvolio is an iron transporter homologous to mammalian DMT1, expressed in the midgut, Malpighian tubules, brain, and testes (Folwell et al., 2006). *Malvolio* mutants exhibit iron deficiency (Bettedi et al., 2011) and ubiquitous RNAi knockdown was previously shown to improve motor performance in a RNAi-based Drosophila FRDA model (Soriano et al., 2016). We first used the ubiquitous da-GAL4 driver to confirm the efficiency of the RNAi knockdown. This led to a 78% decrease of transcript levels in third instar larvae (Fig. 2D). Then, we used the Mex-GAL4 driver to downregulate *Malvolio* specifically in the gut, in frataxin-deficient flies. We observed a strong improvement of survival in fh-GAAs flies bearing the two Mex-GAL4 and UAS-Mvl RNAi transgenes, compared to flies carrying only the Mex-GAL4 driver (+147 % of mean lifespan) or only the RNAi transgene (+ 90% of mean lifespan) (Fig. 2E). Altogether, these results show that reducing intestinal iron absorption improves the survival of frataxin-deficient flies.

**Figure 2:**
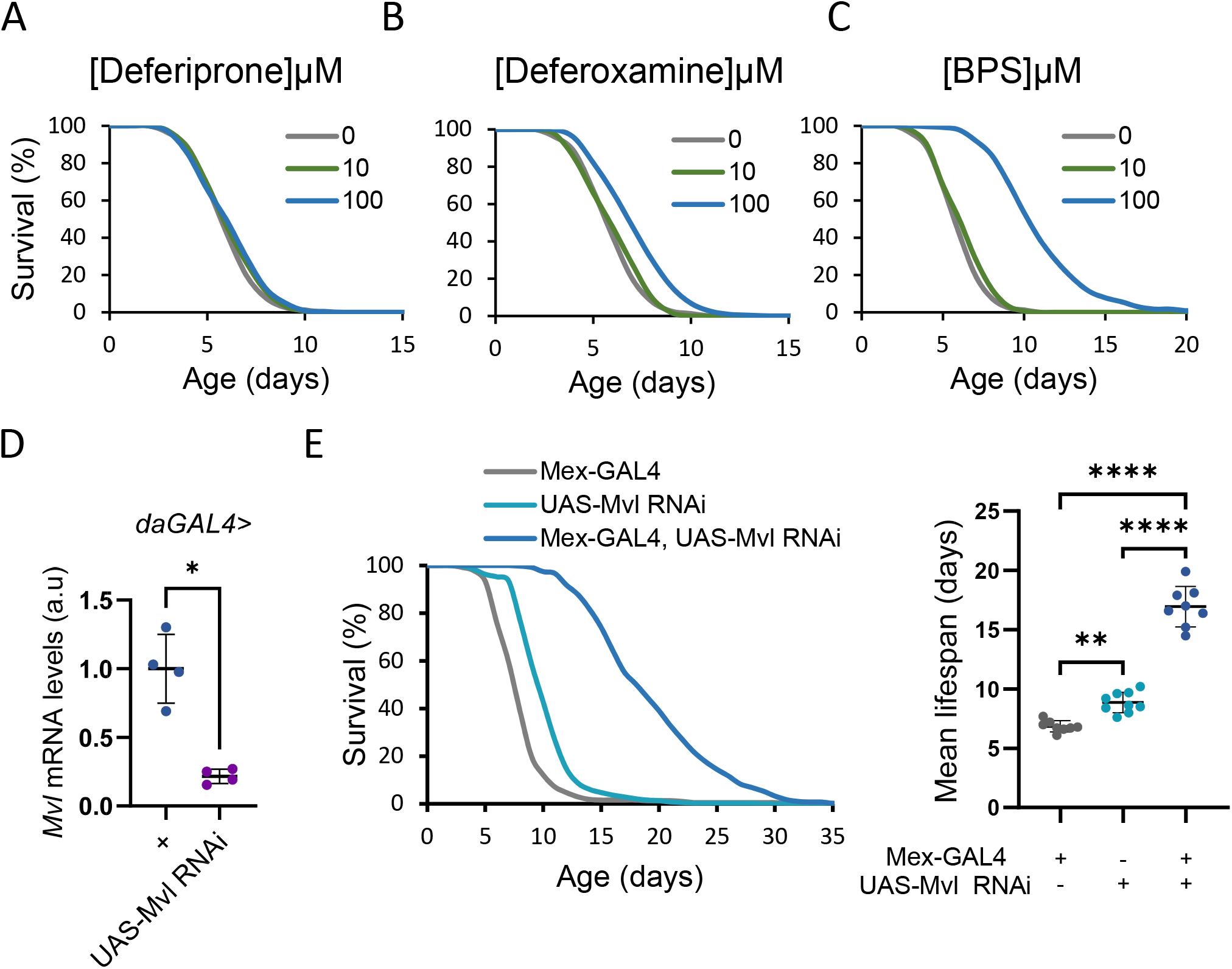
Effects of iron chelators and gut-specific Malvolio KD on the survival of frataxin-deficient flies. A. Survival curves of fh-GAAs male flies treated during development with 10 µM (n=150, p=0,34) or 100 µM (n=219, p=0,12) of deferiprone. B. Survival curves of fh-GAAs male flies treated during development with 10 µM (n=223, p=0,44) or 100 µM (n=235, p=5, 1.10^-11^) of deferoxamine. C. Survival curves of fh-GAAs male flies treated during development or with 10 µM (n=142, p=0,14) or 100 µM (n=230, p=2.10^-71^) of BPS. A-C. Survival curves of untreated fh-GAAs male flies were used as controls (0, n=156). The p-values were obtained in logrank tests using the survival curve of these control flies as the reference. D. Relative quantification of *Mvl* transcripts following RNAi-induced Mvl KD with the ubiquitous da-GAL4 driver. Dots correspond to independent biological samples each obtained from 20 third instar male larvae. Significant difference is indicated (two-sided non-parametric Mann-Whitney tests). E. Survival curves and mean lifespan of *fhGAAs; Mex-GAL4* ; *UAS-dicer2* (Mex-GAL4, n=239), *fhGAAs; UAS-Mvl RNAi* ; *UAS-dicer2* (UAS-Mvl RNAi, n=265), *fhGAAs; Mex-GAL4, UAS-Mvl RNAi* ; *UAS-dicer2* (Mex-GAL4, UAS-Mvl RNAi, n= 244). The UAS-dicer2 transgene was used here to enhance RNAi efficiency. For each condition, flies were housed in tubes by groups of mainly 28 to 30 individuals. Each dot in corresponds to one tube, with mean ± s.d. Significant differences are indicated (One-way Anova followed by Post-hoc Tukey analysis for multiple comparisons). * P < 0.05, ** P < 0.01, *** P <0.0001.

### 3.3 BPS treatment limits iron-induced developmental lethality, reduces CNS iron overload and improves locomotor function but fails to prevent cardiac dilatation

To further link BPS protective effects in fh-GAAs flies to its iron chelation properties, we evaluated the effects of BPS on iron-induced developmental toxicity. Hypersensitivity to iron supplementation was previously reported in the drosophila *fh*^*1*^ mutant and RNAi-induced models (Anderson et al., 2005; Chen et al., 2016; Navarro et al., 2015). Here, fh-GAAs flies were highly hypersensitive to dietary iron : when *w*^*1118*^ control flies developed normally on a medium supplemented with up to 5mg/ml of ferric ammonium citrate (FAC), less than 5% of fh-GAAs flies reached the adult stage with a 25 µg/ml FAC supplementation. Thus, we compared the developmental viability of fh-GAAs flies submitted to 10 to 20µg/ml FAC treatments, with or without 100µM BPS. Only 26% of fh-GAAs flies reached the adult stage on a 10µg/ml FAC medium, compared to 80% when BPS was also added (Fig. 3A). On a 15 µg/ml FAC medium, 8% of fh-GAAs flies were viable, compared to 46% with FAC and BPS. We failed to detect improved viability on a 20 µg/ml FAC medium, which might be due to exceeded BPS iron-chelation capacities at this FAC concentration. We also evaluated the effects of copper supplementation on viability. On a medium supplemented with 1mM copper, less than 3% of fh-GAAs flies reached the adult stage (Fig. 3A), when control flies were unaffected. We evaluated three lower copper concentrations but could not detect any differences in viability following addition of BPS.Therefore, BPS treatment partially rescued iron-induced but not copper-induced developmental lethality, further supporting that BPS protective effects in standard rearing conditions are iron-dependent. Next, we assessed BPS effects on larval brain iron accumulation, which was strongly reduced in fh-GAAs flies when BPS was added to the medium (Fig. 3B). We subsequently monitored cardiac function. As previously described (Jullian et al., 2024), fh-GAAs flies exhibited cardiac dilatation, with systolic and diastolic diameters increasing by 83% and 50% respectively. BPS treatment did not modify the systolic diameters of fh-GAAs and *w*^*1118*^ flies, nor the diastolic diameters of *w*^*1118*^ flies (Fig. 3C). We even observed a slight increase of 14% of fh-GAAs diastolic diameter following BPS treatment. Thus, BPS failed to rescue cardiac dilatation in frataxin-deficient flies. Finally, we evaluated the locomotor function. While four days old fh-GAAs flies showed a marked decrease of spontaneous locomotor function, BPS treatment increased their daily activity by 76% (Fig. 3D). Following activity profiles on a 24 hour period, we observed that the treatment particularly improved the onset and amplitude of the two activity peaks in the morning and the evening anticipating lights on and off (Fig. 3E). Thus, a pharmacological treatment limiting iron absorption in the gut efficiently reduces iron accumulation in the larval CNS and improves adult spontaneous activity.

**Figure 3:**
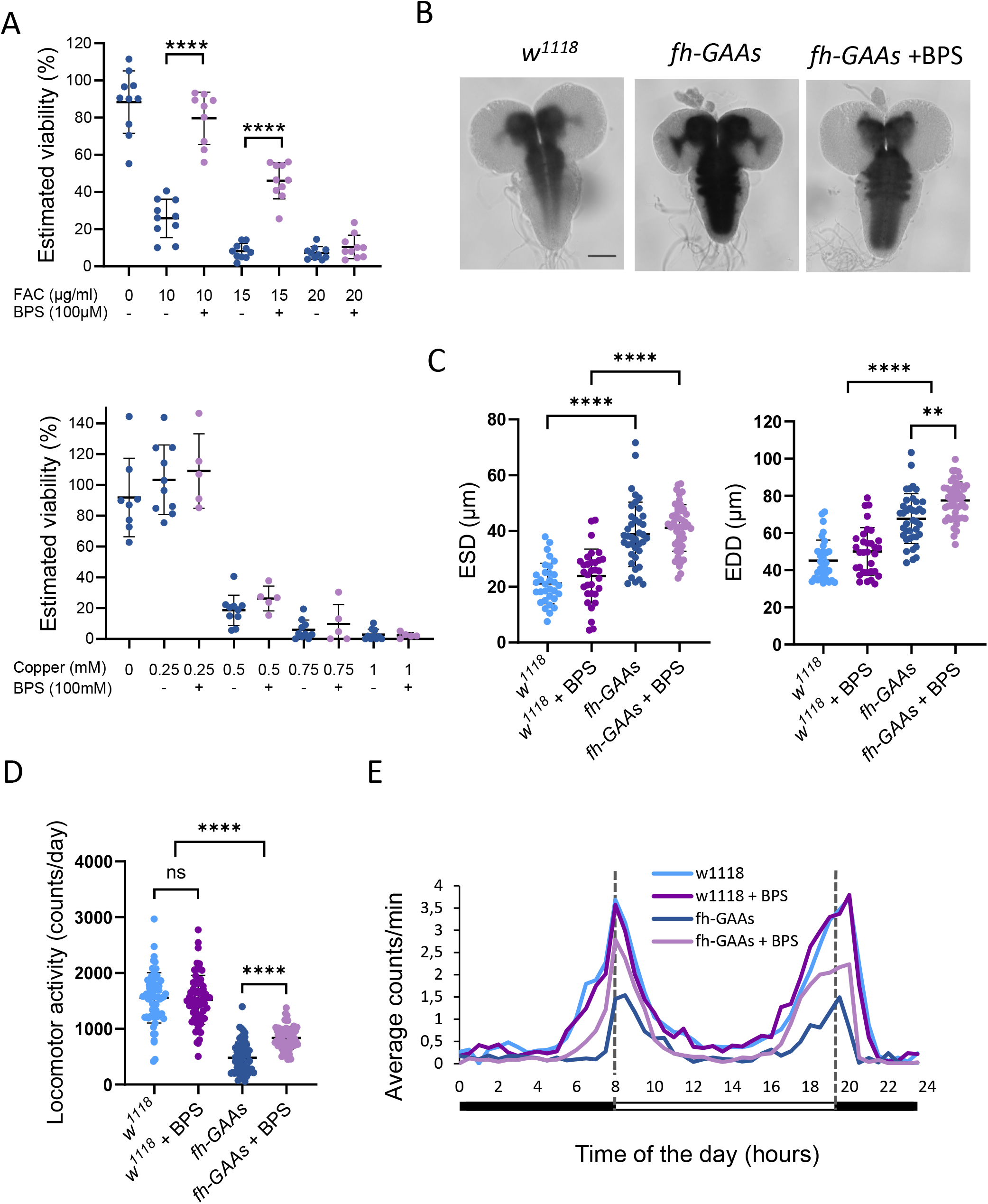
Effects of BPS treatment on dietary iron toxicity, cardiac dilation and locomotor activity. A. Estimated viability of fh-GAAs male flies treated with BPS and ferric ammonium citrate (upper panel) or copper (lower panel). Significant differences between BPS-treated and untreated flies for a given metal concentration are indicated. B. Iron staining of *w*^*111*8^ control and fh-GAAs CNS of third instar male larvae, untreated or treated with 100 µM BPS (scale bar : 100 µm). C. End-Systolic (ESD, μm) and End-Diastolic Diameters (EDD, μm). Cardiac imaging was performed on 7 days old untreated (n=32) or BPS-treated (n=32) *w*^*1118*^ ; *Hand GS>mitoGFP*, and on untreated (n=39) or BPS-treated (n=49) *fh-GAAs* ; *HandGS>mitoGFP* male flies. E-F. Spontaneous locomotor activity of 4 days old untreated and BPS-treated *w*^*1118*^ and *fh-GAAs* adult male flies (n=64 per condition). D. number of counts per day. Each dot corresponds to a single fly. E. Daily activity profiles given as the average counts per minute (mean value over 30 successive minutes) for each condition, in function of the time of the day in hours. The 12h-12h light dark cycle is indicated (lights on and off at respectively 8 am and 8 pm). Flies were treated with 100 µM BPS during development and adulthood. Significant differences are indicated: ** P<0.01,*** P<0.001, **** P<0.0001 (Two-way Anova followed by Post-hoc Tukey analysis for multiple comparisons).

### 3.4 BPS treatment restores a normal larval CNS size and improves glial phenotypes

While assessing iron accumulation in the CNS in previous experiments, we noticed that fh-GAAs CNS were smaller compared to controls. To confirm this, we dissected additional larvae and confirmed that brains and ventral nerve cords of wandering fh-GAAs third instar larvae exhibited reduced size compared to control larvae. This was fully prevented by BPS-feeding (Fig. 4A). Next, we performed immunostaining to visualize neuronal and glial nuclei using antibodies recognizing respectively the elav and repo proteins. We focused our analysis on the ventral nerve cord (VNC) of third instar wandering larvae (Fig. 4B-E), since we observed a significant iron accumulation in this region. We observed abnormal profiles of glial nuclei, with reduced size and flattened shape reflected by an increased aspect ratio (Fig. 4B). We also measured the ratio between the number of glial and neuronal nuclei, which was decreased in fh-GAAs (Fig. 4C). BPS treatment fully prevented these glial abnormalities (Fig. 4B-E). Altogether, these observations strongly suggest that frataxin deficiency triggers iron-dependent cell death early and preferentially in glial cells, a process that can be prevented by dietary iron restriction.

**Figure 4:**
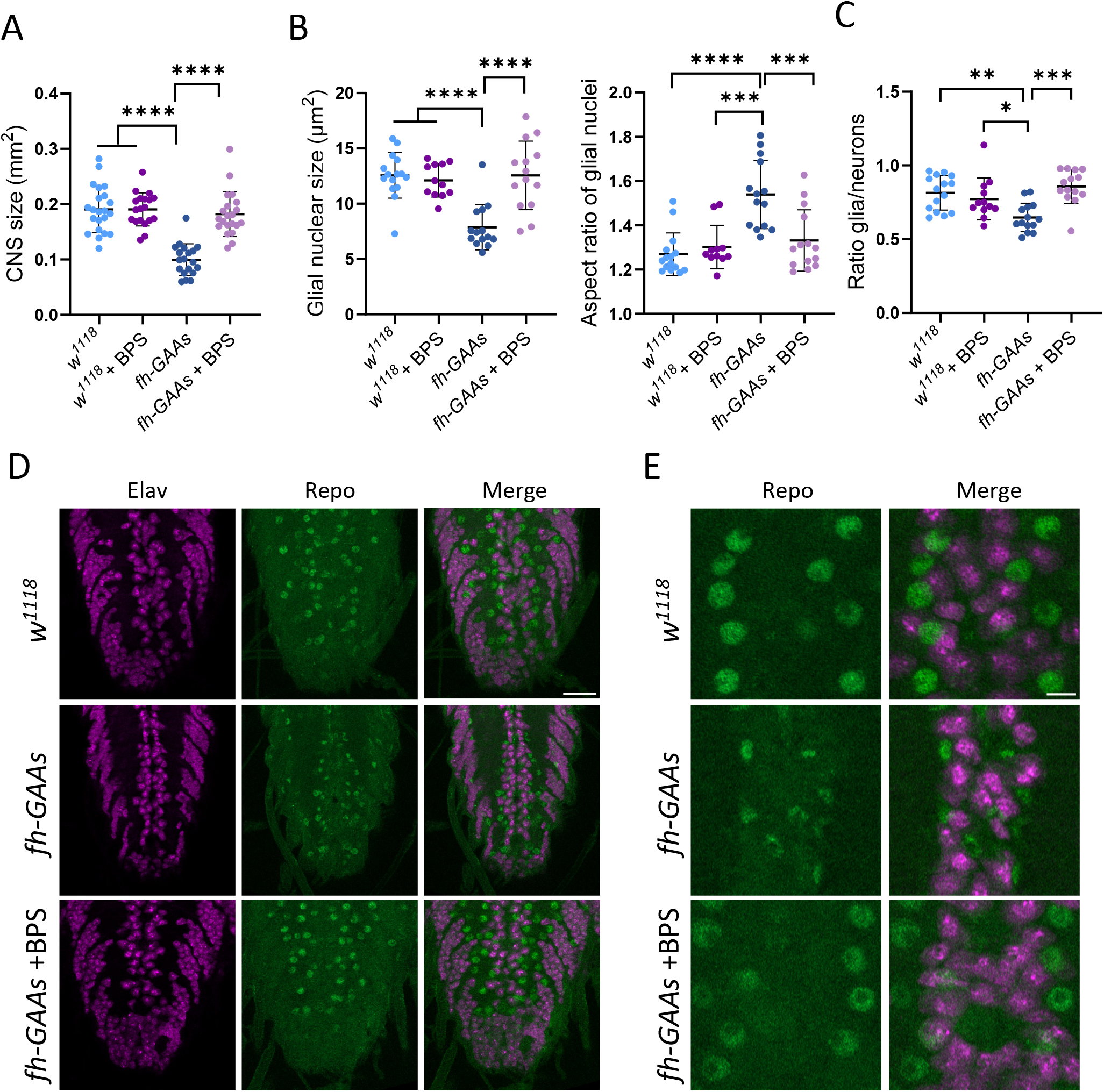
Effects of BPS treatment on larval brain histology. A. CNS size measured as the total surface of the brain and ventral nerve cords. B-E. Immuno staining and evaluation of glial phenotypes on larval ventral nerve cords. B. quantification of glial nuclear size and aspect ratio. Each dot on B-C corresponds to the mean value for a single fly. C. Glia/neuron ratios calculated as the ratio of the number of nuclei detected in the region including the midline and surrounding the two neuropiles. D-E. Immuno staining on larval ventral nerve cords with anti-repo and anti-elav antibodies staining respectively glial and neuronal nuclei. Scale bars : 20 µm (D) and 5µm (E). P<0.05, ** P<0.01, *** P<0.001, **** P<0.0001 (Two-way Anova followed by Post-hoc Tukey analysis for multiple comparisons).

## 4. Discussion

Friedreich ataxia, the most frequent autosomal recessive hereditary ataxia in Caucasians, remains currently incurable. While Omaveloxolone, an NRF2 activator, recently emerged as the only approved treatment for FRDA (Lynch et al., 2021), it only slows down the progression of the disease and is associated with mild adverse effects, such as aminotransferase elevations (Gunther et al., 2025). Consequently, there is an urgent need for complementary approaches that might ultimately lead to combinatorial therapies. In this study, we investigated the impact of dietary iron restriction on the dysfunctions induced by frataxin-deficiency. Our GAA-based *Drosophila* model was proved to be particularly suitable for this objective : by inducing a systemic yet partial depletion of frataxin, it more closely mimics the clinical situation of the patients compared to complete loss of function and tissue- or cell type-specific models. Furthermore, it allows for the *in vivo* assessment of a broad spectrum of frataxin-related phenotypes, enables the use of genetic tools to modulate intestinal iron absorption and the fast evaluation of dietary interventions.

As a first step, we characterized iron dyshomeostasis within our model. In control larvae, the Prussian blue staining which detects Fe^3+^ was not uniformly distributed throughout the central nervous system. In fh-GAAs flies, iron accumulation largely followed the same endogenous pattern, with strong accumulation in the two hemispheres of the central brain, the neuropile of the ventral nerve cord and symmetric structures consistent with the larval optic neuropiles. Thus, iron accumulates in regions of the central nervous system where it is normally more present. While Fe^3+^ accumulation was previously reported in *fh*^*1*^ mutant, carrying a severe loss of function missense mutation leading to L3-pupal lethality, it exhibited a more diffuse pattern compared to the restrained localization observed here (Chen et al., 2016). This points toward a potential link between the severity of functional frataxin deficits and the spatial extent of iron deposits in the nervous system. Strikingly, no additional sites of iron accumulation were detected in fh-GAAs flies, except in pericardial cells, which are large cells arranged in two rows on both sides of the heart. These cells exhibit nephrocytic functions, filtering the hemolymph and exhibiting high levels of ferritin (Mehta et al., 2009). Iron accumulation in these cells may suggest a sequestration mechanism to counteract elevated circulating and systemic iron. However, our systemic data do not support this hypothesis, as whole-body ferrozine assays—detecting both ferric and ferrous iron—revealed no significant increase in total iron levels. Furthermore, we found no major systemic changes in the expression of key regulators of iron homeostasis. Interestingly, tissue-specific extents of iron accumulation were also recently described in mouse with partial systemic frataxin deficiency (Pazos-Gil et al., 2026). Better understanding the underlying mechanisms of these tissue specificities clearly requires further investigation

Considering the unsolved debate in the FRDA field about the involvement of iron accumulation in the progression of the disease and the relevance of using iron chelators, we aimed to evaluate this approach on our model. Deferoxamine and deferiprone are iron chelators used to treat iron overload across various clinical conditions (Entezari et al., 2022). Deferoxamine, which possesses a high affinity for ferric iron, has been the standard treatment in patients with thalassemia for several decades, with intravenous or subcutaneous administration due to its poor oral bioavailability (Xia et al., 2013). Deferiprone is administered orally and capable of crossing the blood-brain barrier and mitochondrial membranes, properties that led to its evaluation in patients with FRDA. While a slight decrease of cardiac hypertrophy was observed, the treatment was ultimately disregarded due to significant adverse effects such as worsening of tremor (Arpa et al., 2014; Elincx-Benizri et al., 2016; Pandolfo et al., 2014). BPS is a membrane-impermeable iron chelator with high affinity for ferrous iron, currently used in cell cultures to chelate iron in the medium. In Drosophila, BPS efficiently prevents iron absorption upon dietary administration, by capturing and precipitating iron into granules, particularly in a small region of the anterior midgut presumed to be a major site of iron absorption (Hernandez-Gallardo et al., 2024). We observed striking discrepancies in the ability of these three iron chelators, administered in the diet, to extend the survival of fh-GAAs flies. While deferiprone showed no effect, deferoxamine and BPS exhibited weak and strong effects, respectively—with BPS nearly doubling the lifespan of frataxin-deficient flies. These findings highlight that extracellular chelators, by sequestering dietary iron and thereby limiting its intestinal absorption, represent the most effective pharmacological approach in this model. This mechanism was further validated by a genetic approach targeting the Malvolio iron transporter. We show here that its targeted downregulation specifically in the gut is as efficient as BPS treatment in extending adult survival. Notably, these two interventions— BPS treatment and gut-specific *Malvolio* knockdown—emerged as the most potent strategies among all pharmacological and genetic conditions we have previously tested in *fh-GAA* models, except the full rescue through UAS-GAL4 driven ubiquitous expression of Drosophila or human frataxin (Jullian et al., 2024; Russi et al., 2020; Want et al., 2025). Besides survival improvement, treatment with BPS also improved adult locomotor activity, limited iron-induced developmental lethality and rescued the reduced brain size. In contrast, cardiac function was not improved, which is consistent with the absence of marked iron accumulation in cardiomyocytes. This led us to conclude that cardiac dilation in fh-GAAs flies is not a direct consequence of cardiac iron overload. This observation is in agreement with murine models, in which iron accumulation occurs secondary to initial cardiac dysfunction (Puccio et al., 2001).

A major finding of our study is that glial cells are preferentially affected in the CNS of frataxin-deficient flies, in an iron-dependent manner. Although FRDA has traditionally been viewed as a neuron-centered disease, accumulating evidence indicates that glial cells also contribute to disease pathogenesis. In Drosophila, glia-specific frataxin silencing results in locomotor impairment and reduced lifespan (Navarro et al., 2010). Moreover, frataxin-deficient human astrocytes exhibit mitochondrial dysfunction and promote neuronal damage, highlighting the importance of non-cell-autonomous mechanisms in FRDA (Loria and Diaz-Nido, 2015). Neuropathological studies have also revealed marked abnormalities of satellite glial cells in dorsal root ganglia from FRDA patients (Koeppen et al., 2016), while Schwann cells display pronounced vulnerability to frataxin deficiency, including cellular dysfunction and cell death (Lu et al., 2009). Our model provided the opportunity to directly compare neuronal and glial populations carrying the same *fh* mutant allele within the same tissue environment. At the developmental stage examined here, glial cells exhibited substantially more severe alterations than neurons, supporting the emerging view that glial dysfunction represents an early pathogenic event in FRDA. The complete rescue of this phenotype by BPS further indicates that iron-dependent mechanisms contribute to this early vulnerability of glial cells. Accumulating evidence indicates that ferroptotic pathways are active in FRDA models and omaveloxolone, the only approved drug for FRDA, was recently shown to improve ferroptotic markers in DRG neurons (Portillo-Carrasquer et al., 2026). Here, a ferroptosis-like mechanism is strongly suggested by the fact that glial abnormalities are rescued by BPS treatment. However, classic ferroptosis is characterized by a nucleus that remains morphologically intact at least during early stages (Xie et al., 2016). In this respect, the small, flattened phenotype of the glial nuclei observed in our model is atypical but might reflect late-stage architectural collapse and nuclear envelope distortion preceding cell death. Whether the glial abnormalities observed in our model are linked to ferroptotic events remains to be confirmed using additional ferroptotic markers. Interestingly, glial cells were shown to be the most iron-rich cells in the brain in rats (Reinert et al., 2019). In this study, the resolution of our iron staining method did not allow us to determine whether iron accumulation was more pronounced in glial cells than in neurons in control and frataxin-deficient flies. Exploring this distinction will be crucial as it could reveal a potential mechanism to explain the specific glial vulnerability to frataxin deficiency.

Finally, we identified intestinal iron uptake as an important determinant of disease severity in our Drosophila model of frataxin deficiency. These effects were observed in the absence of systemic iron overload, suggesting that disease severity is influenced not only by local iron dysregulation but also by the dynamics of iron entry into the organism. While these findings do not directly support dietary iron limitation as a therapeutic recommendation for patients, they highlight systemic iron flux as a potential contributor to FRDA pathophysiology. Further studies in mammalian models will be required to determine whether modulation of intestinal iron absorption and dietary iron can be safely and effectively exploited in a clinically relevant context.

## Author contribution

**Ema Turki :** Conceptualization, Methodology, Investigation, Validation, Formal analysis, Visualisation, Writing-Review and Editing. **Estelle Jullian :** Conceptualization, Methodology, Investigation, Validation, Formal analysis. **Pierre Delamotte :** Methodology, Investigation, Formal analysis. **Anne Filipe :** Investigation, Validation, Writing-Review and Editing. **Laura Tixier-Cardoso :** Investigation. **Sandrine Middendorp :** Writing-Review and Editing. **Elodie Martin :** Writing-Review and Editing. **Véronique Monnier :** Conceptualization, Methodology, Investigation, Validation, Formal analysis, Supervision, Project administration, Funding acquisation, Writing-Original Draft, Review and Editing.

## Acknowledgements

We thank Hervé Tricoire for his useful comments on the manuscript. We acknowledge the ImagoSeine core facility of the Institut Jacques Monod, member of the France BioImaging infrastructure (https://ror.org/01y7vt929) supported by the French National Research Agency (ANR-24-INBS-0005 FBI BIOGEN) and GIS-IBiSA.

