## Supplementary Fig. 1 and Table S1 for "Limiting intestinal iron absorption rescues glial defects and extends lifespan in a Drosophila model of Friedreich’s ataxia"

### Supplementary Data

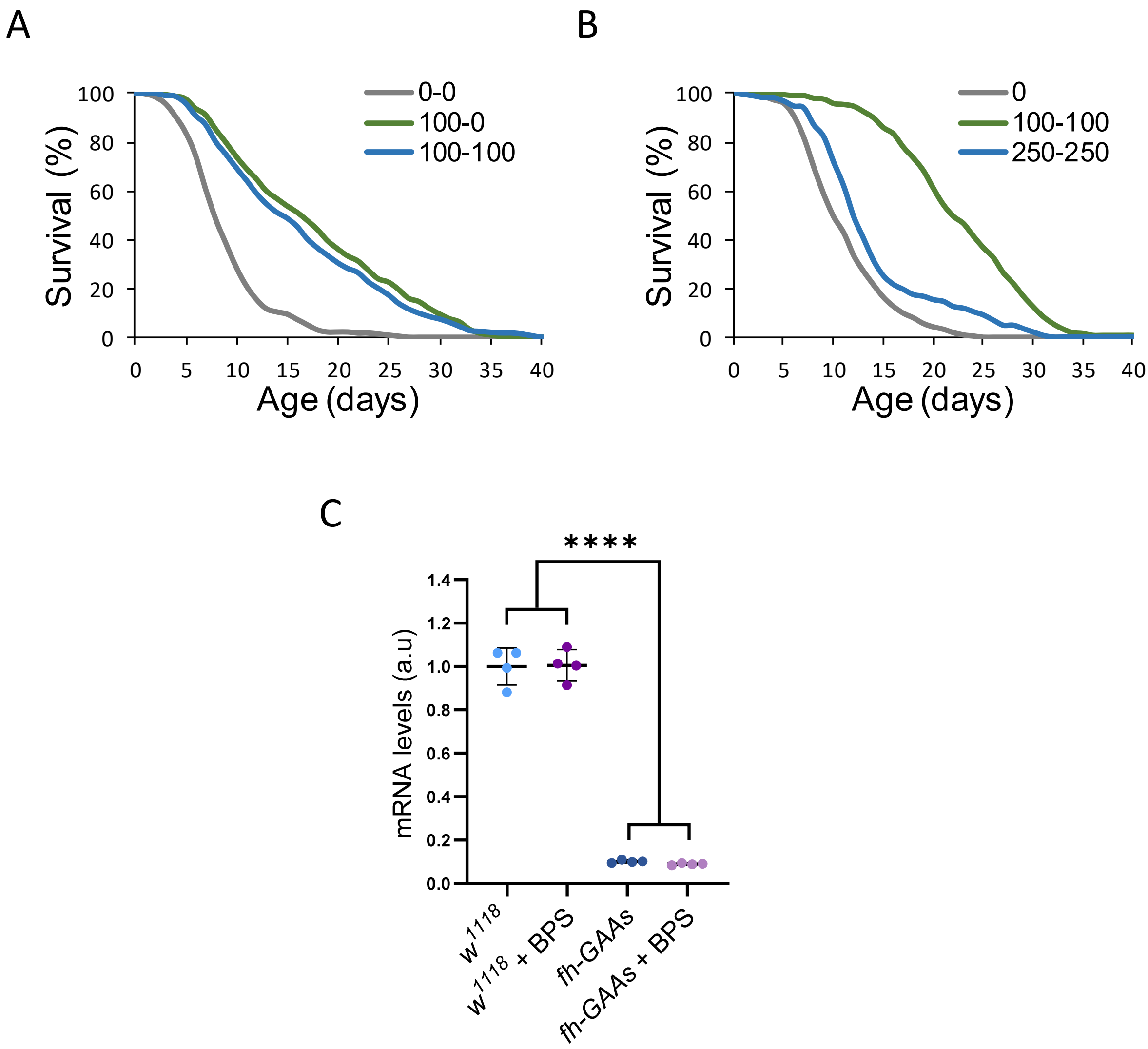


**Figure S1 : Effects of BPS treatments on adult survival and *fh* expression**

A. Survival curves of fh-GAAs male flies untreated (n=232), treated only during development (n=210) or continuously (n=295) with 100 µM of BPS. No significant differences were detected between the two treatments (p=0,16). B. Survival curves of fh-GAAs male flies untreated (n=215), or treated continuously with 100 µM (n=273) or 250 µM (n=144) of BPS. The 250 µM BPS treatment significantly increased survival compared to untreated flies ($p=2,1.10^{-6}$) but were significantly less efficient compared to the 100 µM treatement ($p=3,5.10^{-30}$). The p-values were obtained in logrank tests C. Quantification of *fh* transcript levels in control $w^{1118}$ and fh-GAAs male third instar larvae, untreated or treated with BPS. Dots correspond to independent biological samples each obtained from 20 individuals. **** P<0.0001 (Two-way Anova followed by Post-hoc Tukey analysis for multiple comparisons).

| Gene | CG number | Sequence |
|---|---|---|
| *Fer1HCH* | *CG2216* | TAATTGCTAGCCTGCTCCTGT |
| | | ATCTCCATAGGCCTGGGC |
| *Fer2LCH* | *CG1469* | GCACTCGCTCTCTTTGCGT |
| | | GGTGATTACAGTGTTCTGGCAAT |
| *Mfrn* | *CG4963* | AGCACACTCGCTATACTTTGC |
| | | GTTGAGGTTTCTCACCGATGT |
| *tsf1* | *CG6186* | CGCACGTCCTGAAGGTGTC |
| | | GACTGCGTGAAGAACTCGGA |
| *rp49* | *CG7939* | CCGCTTCAAGGGACAGTATCT |
| | | CACGTTGTGCACCAGGAACTT |
| *fh* | *CG8971* | ACACCCTGGACGCACTGT |
| | | GTTGATCACATAGGTGCCGTG |
| *Malvolio* | *CG3671* | CATACCCGACGATGATAGCAC |
| | | CGCAATGGACATAAGAAAACCG |

**Table S1 : Primers used for qPCR**
